# FLP-15 modulates the amplitude of body bends during locomotion in *Caenorhabditis elegans*

**DOI:** 10.1101/2025.05.08.652979

**Authors:** Umer Saleem Bhat, H Sharanya, Siju Surendran, Namra Tasnim, Kavita Babu

## Abstract

Locomotion is essential for executing most behaviours. In *Caenorhabditis elegans*. Efficient locomotion is exhibited as a result of the coordination of excitatory and inhibitory signals from the nervous system onto the body-wall muscles. Although neurotransmitters play a vital role in maintaining and executing coordinated movements, neuropeptides have emerged as important players in the regulation and sustenance of locomotory states. In our previous study we explored the role of the neuropeptide FLP-15 in regulating reversal frequency during foraging behaviour in *C. elegans*. We were also interested in exploring other possible locomotory defects in *flp-15* mutant animals. In this work we show that *flp-15* mutants show an increased length of reversals during foraging resulting in defects in maintaining the direction of reversals. Mutants in *flp-15* exhibited a “floral” pattern of reversals as opposed to near linear patterns of reversal in wild-type control animals. We further show that the defect in maintaining the direction of reversals could be due to increased amplitude of the body-bends with *flp-15* mutants showing a large increase in the mean amplitude of body-bends. Our data suggests that FLP-15 partially functions through the G-protein coupled receptor (GPCR), NPR-3, to regulates the amplitude of body-bends. Finally, we show that loss of *flp-15* leads to an increase in the expression of another neuropeptide, NLP-12, whose over expression has been implicated in causing increased amplitude of body-bends allowing us to speculate that the regulation of NLP-12 by FLP-15 may allow for the observed locomotory defects in *flp-15* mutant animals.

## Introduction

Locomotion is a fundamental aspect of behaviour in many organisms, enabling essential activities such as foraging for food, avoiding predators or aversive stimuli, and mating (reviewed in (Domenici *et al*, 2007)). In *Caenorhabditis elegans*, locomotion is crucial for the animal’s survival and reproductive success. This locomotion is characterised by a sinusoidal wave-like motion along the dorsoventral axis. This wave-like motion is generated through a finely tuned coordination of neural and muscular activities (Alfonso *et al*, 1993; McIntire *et al*, 1993a; McIntire *et al*, 1993b; White *et al*, 1976, 1986). The nervous system and musculature of *C. elegans* are arranged in a way that allows for efficient and smooth movement, highlighting the intricate relationship between neural signalling and muscular response (White *et al*., 1986).

The ventral nerve cord in *C. elegans* plays a central role in this locomotion by harbouring the cell bodies of key motor neurons. Among these are the excitatory cholinergic motor neurons, specifically the A and B-type neurons, which are responsible for stimulating muscle contraction (Alfonso *et al*., 1993; Jospin *et al*, 2009). These neurons release the neurotransmitter acetylcholine, which binds to receptors on the muscle cells, causing them to contract and generate movement. This excitation is essential for initiating and maintaining the forward and backward waves of movement that characterizes locomotion in *C elegans* (Jospin *et al*., 2009).

In contrast, the inhibitory GABAergic motor neurons, known as D-type neurons, modulate activity by preventing excessive muscle contractions, thereby ensuring balanced and coordinated movements (Kreyden *et al*, 2020; Liu *et al*, 2020; McIntire *et al*., 1993a; McIntire *et al*., 1993b; Safdie *et al*, 2016). These neurons release gamma-aminobutyric acid (GABA), which inhibits muscle activity on the side opposite to the contraction, allowing for the undulating wave pattern necessary for smooth locomotion. The interplay between excitatory and inhibitory signals ensures that *C. elegans* can move efficiently and respond dynamically to its environment. Therefore, maintaining a critical balance between excitation and inhibition is essential for the execution of normal sinusoidal movement. This balance ensures that motor functions are properly regulated, allowing for smooth and coordinated locomotion (reviewed in (Thapliyal & Babu, 2018)).

Previous studies have highlighted the role of neuropeptides in regulating acetylcholine release at the neuromuscular junction (NMJ) in *C. elegans* (Bhattacharya et al, 2014; Hu et al, 2011; Sieburth et al, 2005). Studies have found that mutants lacking enzymes crucial for neuropeptide maturation showed reduced acetylcholine release. Specifically, *C. elegans* deficient in carboxypeptidase E (encoded by *egl-21*) and proprotein convertase (encoded by *egl-3*) exhibited lower levels of acetylcholine release at the NMJ compared to wild-type animals (Jacob & Kaplan, 2003; Kass *et al*, 2001). These enzymes are essential for processing prepropeptides into mature neuropeptides, suggesting that proper neuropeptide maturation is critical for maintaining normal acetylcholine levels and subsequently normal motor function.

Although multiple studies have explored the role of neuropeptidergic modulation in maintaining the excitation/inhibition (E/I) balance and neurotransmitter release (Banerjee *et al*, 2017; Bhattacharya *et al*., 2014; Flavell *et al*, 2013; Florman & Alkema, 2022; Hu *et al*., 2011; Jacob & Kaplan, 2003; Ji *et al*, 2023; Lim *et al*, 2016; Oranth *et al*, 2018; Pandey *et al*, 2021; Shahi *et al*, 2025; Stawicki *et al*, 2013), the function of a number of neuropeptides involved in this process remains unknown. As an example, neuropeptides like NLP-12 and FLP-18 have so far been shown to play critical roles in feedback mechanisms and homeostatic processes that ensure proper motor control (Bhardwaj *et al*, 2018; Bhattacharya *et al*., 2014; Hu *et al*., 2011; Hu *et al*, 2015; Hums *et al*, 2016; Pandey *et al*., 2021; Ramachandran *et al*, 2021; Stawicki *et al*., 2013; Tao *et al*, 2019).

The neuropeptidergic modulation of neurotransmitter release at the NMJ represents a sophisticated level of regulation that is vital for coordinated movement and behavioural responses (reviewed in (Bhat *et al*, 2021; Li & Kim, 2008)). Understanding the molecular basis of how neuropeptides can significantly influence the release of neurotransmitters and, consequently affect locomotory behaviour will allow for a more complete understanding of the role of neuropeptides in multiple behaviours. These findings have potential implications for understanding similar processes in other organisms and could provide insights into motor control disorders and neurodegenerative diseases where E/I balance is disrupted across phyla (Barbour *et al*, 2024; Bi *et al*, 2020; Ghatak *et al*, 2021; Maestu *et al*, 2021).

Here, we show that the neuropeptide FLP-15 regulates the amplitude of body-bends in *C. elegans*, where *flp-15* mutants show a stark increase in the amplitude of body-bends. Our data suggests two possible pathways for modulation of the amplitude of body-bends in *flp-15* mutants, one via the GPCR receptor NPR-3 expressed in dopaminergic neurons and the other through the modulation of another neuropeptide, NLP-12.

## Materials and Methods

### Strains maintenance

All strains were cultured on nematode growth medium (NGM) plates, inoculated with OP50 *E. coli* bacteria, and maintained under standard conditions at 22°C, following established protocols outlined in previous literature (Brenner, 1974). The N2 Bristol strain served as the standard wild-type (WT) control for comparative analyses across all experimental procedures. A synchronous population of young adult *C. elegans* was obtained using previously established protocol (Porta-de-la-Riva *et al*, 2012). The list of strains used in this study is tabulated in Supplementary Table 1 (S1).

### Molecular biology and transgenes

Transcriptional reporters were generated through molecular cloning of the promoter regions, positioning them upstream to fluorescent protein (GFP) in the pPD95.75 vector or mCherry in the pPD49.26 vector. The length of all promoter sequences was maintained at 2 kb, spanning the genomic region upstream of the start codon for each respective gene and the sequences were extracted from WormBase. Promoters of genes *flp-15, gur-3, npr-3, dat-1 and nlp-12* have been used in this study. The rescue constructs were generated by cloning genomic DNA or cDNA downstream of the respective promoter. All the constructs were microinjected in *C. elegans* to prepare transgenic lines as previously described (Mello & Fire, 1995; Mello *et al*, 1991). The list of primers and plasmids used in this study is tabulated in Supplementary Tables 2 and 3 (S2 and S3) respectively.

### Behavioural assays

All the behavioural assays were performed using single young adult staged *C. elegans* recorded one at a time across several days to ensure consistency and account for variability. Briefly, a well-fed young adult animal was picked with an eye lash pick by using a small drop of halocarbon oil, ensuring a gentle transfer. The *C. elegans* was then placed in an off food NGM plate and allowed to crawl freely for one minute to get rid of any residual food. The *C. elegans* was then gently picked again and placed on the freshly prepared 90 mm NGM (without cholesterol) plate calibrated at room temperature and allowed to move freely for one minute before starting the tracking of the animal. The movement of the animal was recorded for 5 minutes at 8 fps using the MBF Bioscience WormLab imaging system (Roussel *et al*, 2014). The reversal length was calculated for each reversal and averaged over the total number of reversals made over five minutes which was then plotted as single data point. Trajectory of *C. elegans* and amplitude of the body-bends were analysed with the help of WormLab software (MBF Bioscience). The number of *C. elegans* recorded for each genotype was ≥ 20.

### Quantitative PCR

Total RNA was isolated from well fed synchronized adult animals (48h including hatching time). 100ng of total RNA was used to synthesize cDNA using Transcriptor High Fidelity cDNA Synthesis Kit (Roche). Real time PCR was performed using LightCycler 480 SYBR Green I Master (Roche) and LightCycler Instrument (BioRad). The analysis of gene expression was done as previously described using the formula

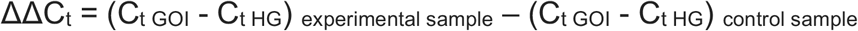

The expression of gene of interest (*nlp-12*) was analysed in WT, *flp-15* and *npr-3* mutant *C. elegans*. The fold increase in gene expression was determined as previously described using formula for fold change = 2^-ΔΔCt^ (Livak & Schmittgen, 2001).

### Statistical analysis

The data was plotted on GraphPad Prism 10 and represented as violin plots and bar graphs according to the distribution of data. Error bars represent SEM. One-way ANOVA with multiple comparison and post-hoc tukey tests was performed to compare the results in all the experiments. The level of significance was set as p≤ 0.05.

## Results

### FLP-15 is required to maintain the direction of reversals

Reorientations including reversals and omega-turns, are critical for food searching behaviours in *Caenorhabditis elegans*. Bhardwaj et al. has shown that the length of reversals is important in determining the extent of the turning angle during reorientation events. Shorter reversals are usually associated with turning angles < 90 degree, while longer reversals are attribute to turning angle between 90-180 degree and in most instances culminate in omega-turns. The study further show that FLP-18 plays a critical role in regulating reversal length in *C. elegans* (Bhardwaj *et al*., 2018). We were interested in further understanding the role of neuropeptides in the regulation of reversal length (Illustrated in Figure 1A). To this end we performed a screen using 22 mutants in neuropeptides to study the role of other neuropeptidergic mutants in regulating reversal length (Bhat *et al*, 2025). We observed a significant increase in the length of reversals in *flp-15* mutants when compared to wild-type (WT) control animals (Figure 1B). Further, this phenotype seen in the mutants was rescued by expressing FLP-15 under its own promoter (Figure 1B). This finding suggests that FLP-15 plays a crucial role in maintaining normal reversal length. While observing the length of reversals, we noted an intriguing phenotype in *flp-15* mutant *C. elegans*. Unlike WT animals that execute reversals in a stereotypic near-straight backward direction, *flp-15* mutants formed a “floral” pattern during reversals (depicted in Figure 1C). This abnormal pattern suggests a significant deviation from the normal movement typically seen in *C. elegans*. The floral pattern observed in *flp-15* indicates a disruption in the coordinated control of body-bends during reversals. This disruption is likely due to an imbalance in the excitatory and inhibitory signals that regulate muscle contractions during movement (reviewed in (Wen *et al*, 2018)).

**Figure 1:**
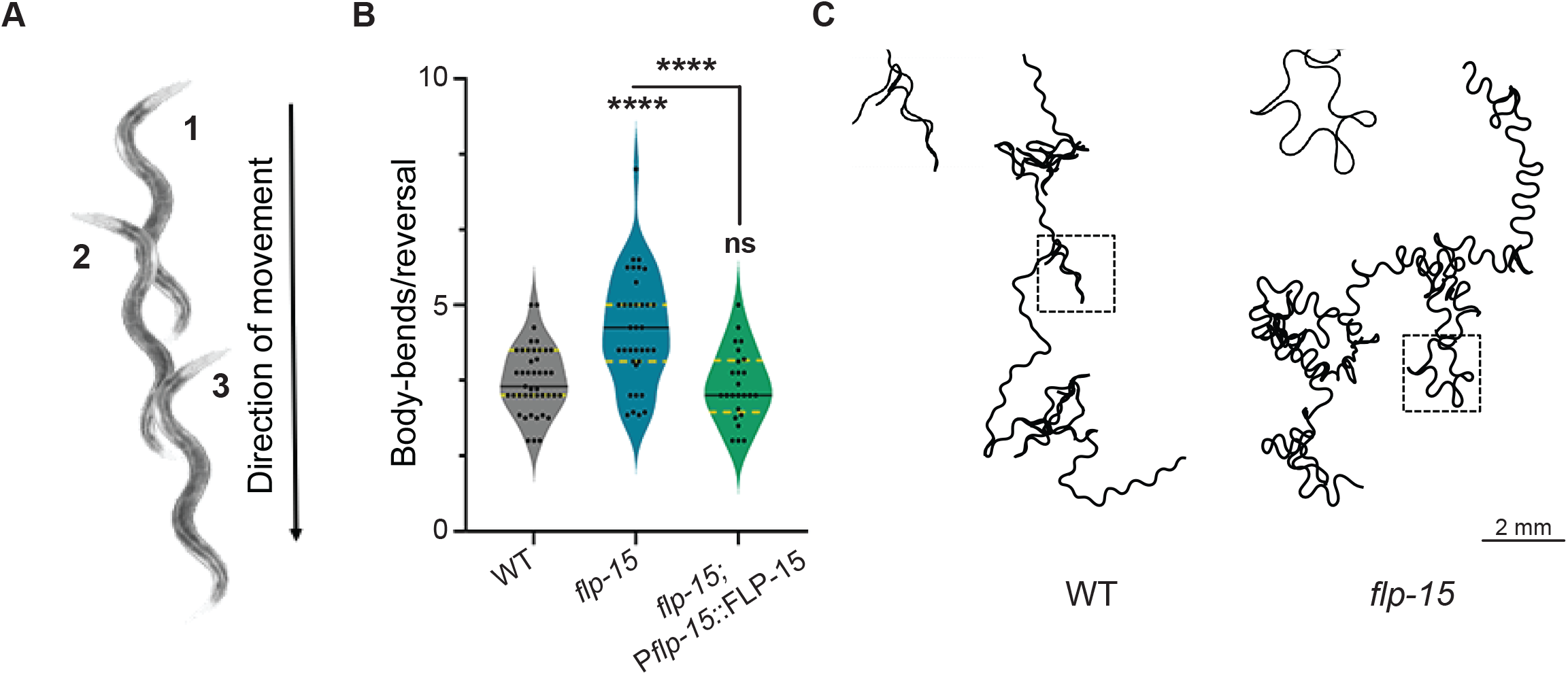
Mutants in *flp-15* show increased body-bends per reversal. (A) Schematic showing body bends (head turning) during reversals, with each number indicating a complete body bend as the head moves in the opposite direction. (B) Quantification of the body bend per reversals. Mutants in *flp-15* show a significant increase in the number of body bends during reversals (reversal length) compared to the wild-type (WT) counterparts. In this plot, each dot corresponds to the number of body bends per reversal from a single *C. elegans*. Statistical significance was determined using One-way ANOVA with the Tukey test and represented by “ns” for not significant and “****” for p< 0.0001. (C) Tracks of *C. elegans* during locomotion. The insets in each track indicate the region containing the dotted square. The insets indicate the linear reversals in WT animals and the floral pattern of reversals in *flp-15* mutants.

To our knowledge there appears to be no previous literature discussing these abnormal reversal patterns in *C. elegans*. Our findings suggest that in addition to length of reversals, maintaining a proper direction of reversal is critical for efficient locomotion and foraging. Proper reversal direction is essential for the effective execution of reorientation movements, which are vital for *C. elegans* to navigate their environment, locate food, and avoid predators. We next examined the locomotion of *flp-15* mutants in some detail and found that *flp-15* is required to maintain the amplitude of body-bends in *C. elegans*.

### FLP-15 regulates the amplitude of body-bends in *Caenorhabditis elegans*

While carefully observing this intriguing defect in the directionality of reversals, we suspected that the ‘floral’ pattern observed in *flp-15* mutants was due to increased amplitude of body-bends. This increased amplitude likely causes exaggerated curvatures while executing reversals, leading to the abnormal reversal patterns seen. To test this hypothesis, we quantified the amplitude of body-bends during forward and reverse movements separately (Illustrated in Figure 2A). We observed that WT animals exhibit higher body-bend amplitude during reversals when compared to making forward movements (Figure 2B). Interestingly in *flp-15* mutant animals there was an increase of amplitude during both forward as well as reverse movements when compared to control *C. elegans*, suggesting a global amplitude defect in these mutants (Figure 2B). To further explore this defect, we quantified the mean amplitude of body-bends across the entire trajectory of the animals (including forward and reverse movements). Our results confirmed that *flp-15* mutants exhibited a significantly large increase in the mean amplitude of body-bends when compared to WT *C. elegans* (Figure 2C). This increased curvature during reversals supports our idea that altered body-bend amplitudes may contribute to the defective directionality in *flp-15* mutants. To further validate our findings, we attempted to rescue this defect by supplementing *flp-15* mutants with FLP-15 cDNA under the control of its endogenous promoter and specifically in the I2 neurons (*gur-3* promoter) where FLP-15 is expressed (Bhat *et al*., 2025). These interventions partially restored the body-bend amplitude in the *flp-15* mutant animals (Figure 2C). These results suggest that the neuropeptide FLP-15 regulates the body-bend amplitude during locomotion to maintain normal locomotion including proper reversal direction. We were next interested in understanding the receptor through which FLP-15 could be functioning to maintain normal body-bends in the animal.

**Figure 2:**
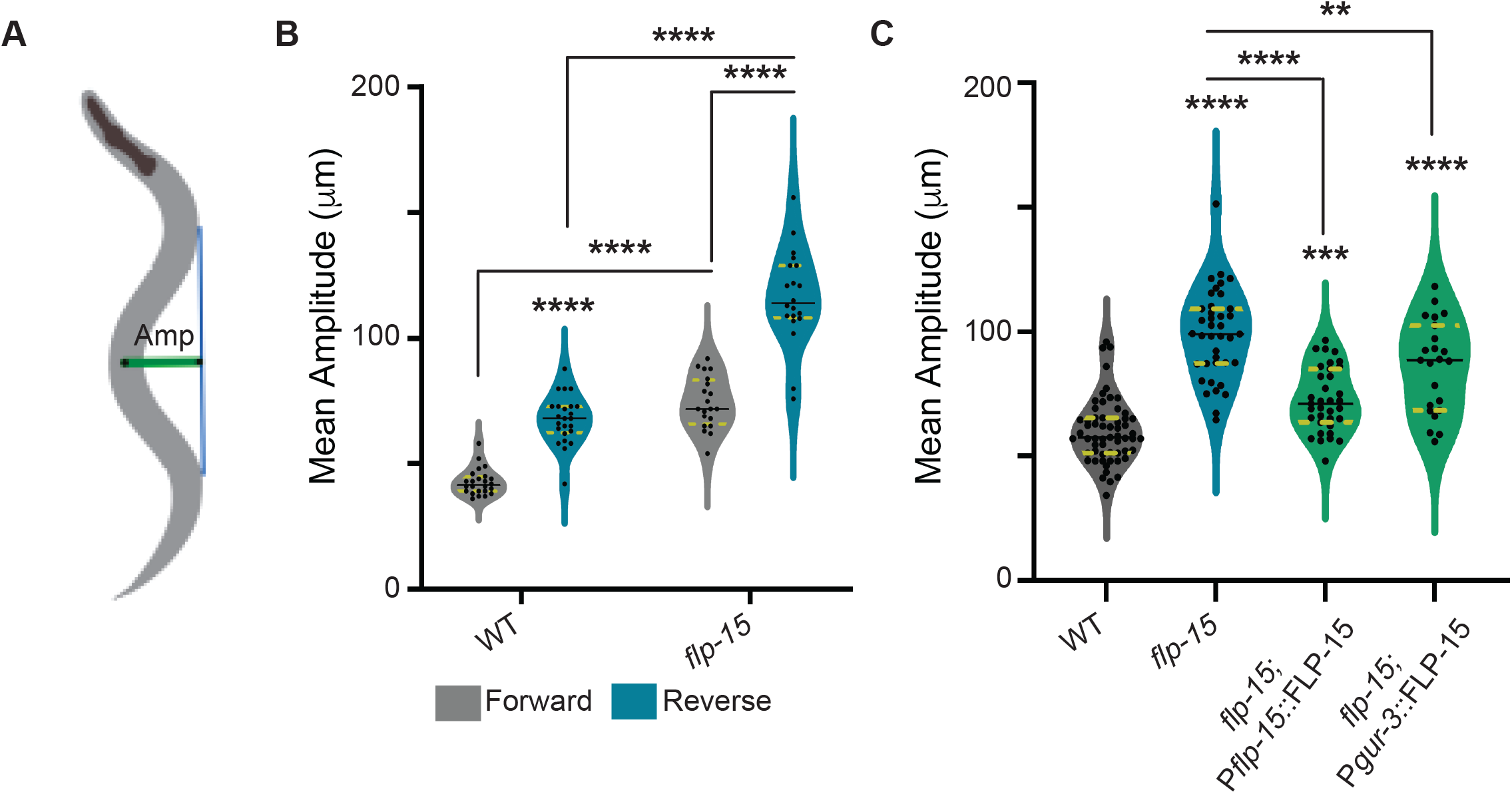
Mutants in *flp-15* show increased amplitude of body-bends per reversal. (A) Schematic showing the measurement of amplitude of body bend during locomotion. The green line indicates the amplitude, and the blue line indicates the baseline of the sinusoidal wave during the movement. The mean of the amplitude is calculated for 5 minutes per *C. elegans* and averaged over multiple animals. (B) Quantification of the mean amplitude during forward and reversal locomotion. Mutant in *flp-15* show a significant increase in the mean amplitude during both forward and reversal locomotion compared to their WT counterparts. In this plot, each dot corresponds to the mean amplitude from a single animal. (C) Quantification of the mean amplitude during the entire trajectory of the animal. Mutants in *flp-15* displayed global defects in the amplitude of body-bends throughout the entire trajectory. The defect was largely rescued by supplementing *flp-15* extrachromosomally under the control of its endogenous promoter and partially by expressing *flp-15* under the I2 neuron-specific promoter (P*gur-3*). Statistical significance was determined for B and C using One-way ANOVA with the Tukey test and represented by “**”, “***’’ or “****” for p-values less than 0.01, 0.001, or 0.0001, respectively.

### FLP-15 regulates the amplitude of body-bends partially through the NPR-3 receptor

Previous work has implicated NPR-3 as a putative receptor for FLP-15 (Beets *et al*, 2023). Moreover, our recent study suggests that FLP-15 functions through NPR-3 to regulate the frequency of reversals during foraging (Bhat *et al*., 2025). We tested whether FLP-15 functions through NPR-3 in regulating the amplitude of body-bends. Our results indicated that *npr-3* mutants show a significant increase in the amplitude of body-bends as compared to WT animals, but this defect is not as severe as that seen in *flp-15* mutants (Figure 3A). These data suggest that NPR-3 may be a receptor involved and likely not the only receptor that binds to FLP-15 to regulate the amplitude of body-bends. We next went on to test the body-bend amplitude in *flp-15; npr-3* double mutant animals and found that the double mutants showed a phenotype similar to that seen in *flp-15* single mutants (non-additive), indicating that *flp-15* and *npr-3* may be functioning in the same pathway to regulate the amplitude of body-bends (Figure 3A). We next overexpressed FLP-15 under its own promoter in *flp-15* and *npr-3* mutant *C. elegans*.

**Figure 3:**
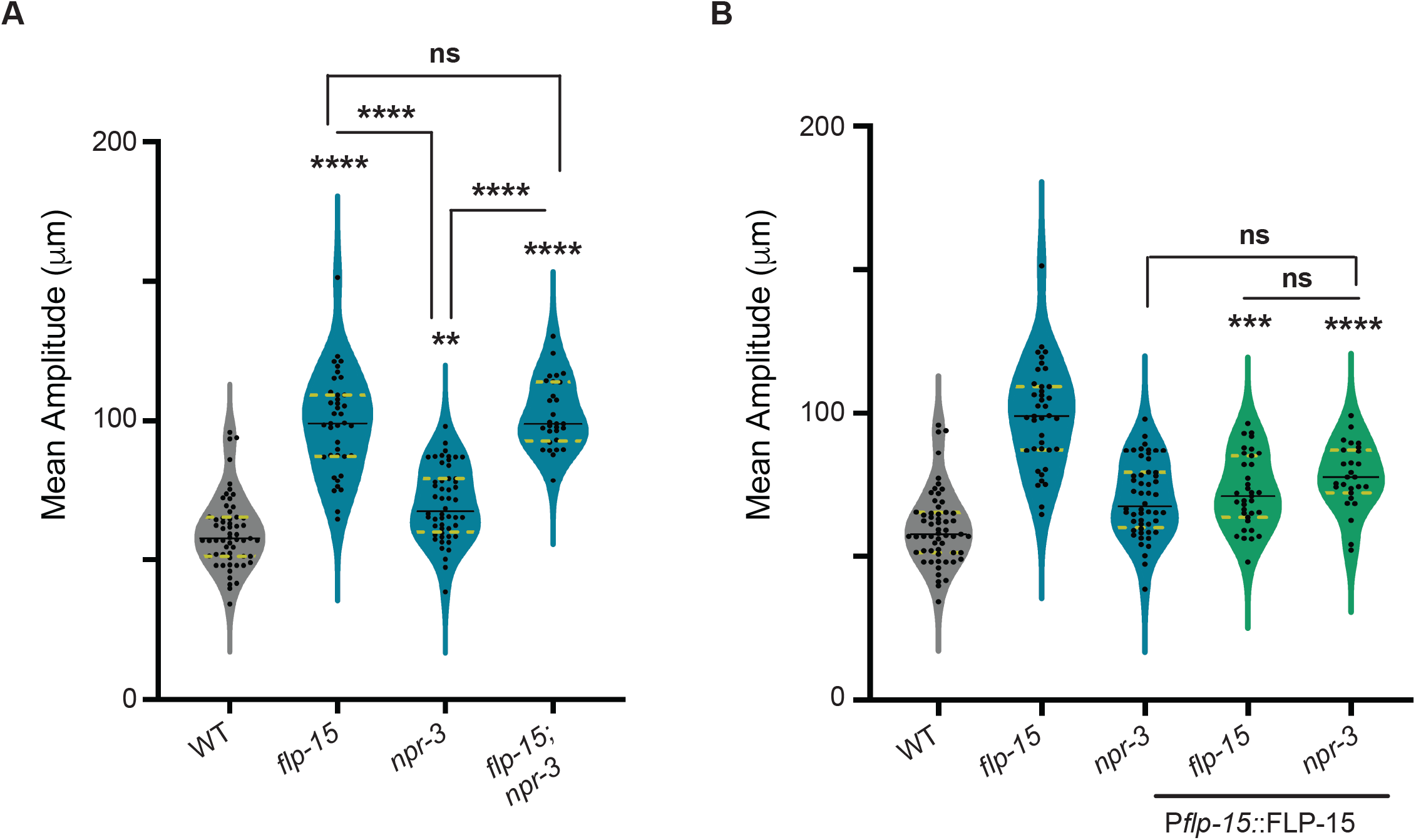
FLP-15 functions partially through NPR-3 to maintain the amplitude of body-bends. (A) Quantification of mean amplitude. The mutants in *flp-15, npr-3 and flp-15; npr-3* double mutants show defects in the amplitude of body-bends during locomotion. This indicates that *flp-15* may function partially through *npr-3*. (B) Overexpression of the *flp-15* gene in the *npr-3* mutant background does not alleviate the defects in the amplitude of body-bends, indicating again that *flp-15* may function partially through *npr-3* to regulate the amplitude of the body-bends. Statistical significance in A and B was determined using One-way ANOVA with the Tukey test and represented by “ns” for not significant, “**”, “***’’ or “****” for p-values less than 0.01, 0.001, or 0.0001, respectively+

We found that *flp-15* mutants show a partial rescue while *npr-3* mutants do not show any change in the increased amplitude of body-bends phenotype (Figure 3B). These data suggest that overexpressing FLP-15 in the absence of *npr-3* does not circumvent the *npr-3* mutant phenotype, again indicating that FLP-15 functions through NPR-3.

We were next interested in studying the site of action of NPR-3 for its role in regulating the amplitude of body-bends in *C. elegans*.

### FLP-15 regulates the amplitude of body-bends partially through NPR-3 expressed in dopaminergic neurons

To further explore the downstream neurons where NPR-3 may be functioning, we conducted neuron-specific rescue experiments in *npr-3* mutant *C. elegans*. Our previous study has suggested that NPR-3 is expressed in ADE and CEP dopaminergic neurons in the head and the DVA neuron in the tail (Bhat *et al*., 2025). We first supplemented the *npr-3* mutants with an extrachromosomal array of wild type NPR-3 genomic DNA (gDNA) under the control of its endogenous promoter. We were able to completely rescue the increased amplitude of body-bends in these animals (Figure 4A). We next expressed the NPR-3 gDNA specifically in dopaminergic neurons using the *dat-1* promoter (Nass *et al*, 2002; Nass *et al*, 2001) or DVA neurons using the *nlp-12* promoter (Bhattacharya *et al*., 2014; Hu *et al*., 2011), in *npr-3* mutants. Our results showed that we were able to partially rescues the defect in the amplitude of body-bend when NPR-3 was supplemented in dopaminergic neurons, however we did not find a significant rescue when NPR-3 was expressed in DVA neurons (Figure 4A). We next asked if dopamine could be involved in this behavioural output. To this end we analysed *cat-2* mutant *C. elegans* that lack the dopamine synthesis enzyme (Sanyal *et al*, 2004). We saw that *cat-2* mutants showed a small increase in amplitude of body-bends similar to but to a lower degree than that seen in *npr-3* mutants (Figure 4B). These data suggest that dopamine signalling through NPR-3 may be involved in regulating the amplitude of body-bends.

**Figure 4:**
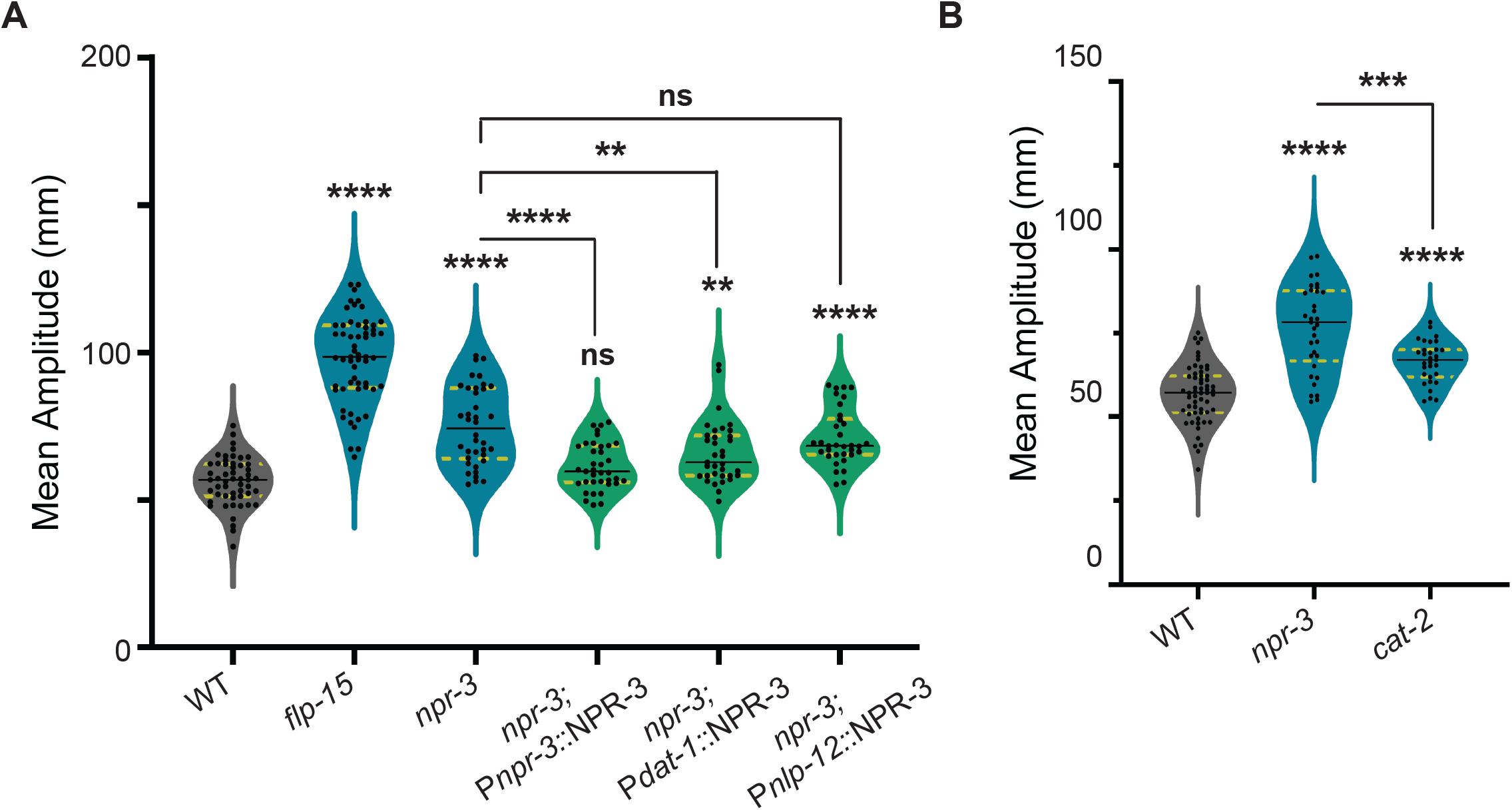
NPR-3 functions in dopaminergic neurons to allow for normal amplitude of body-bends. (A) Quantification of mean amplitude of body bend. The data shows that supplementing *npr-3* gene under the control of its endogenous promoter in the *npr-3* mutant background completely rescues the increased body-bend amplitude phenotype. Additionally, the phenotype is partially rescued when *npr-3* gene is supplemented under the control of the *dat-1* promoter, that drives expression of NPR-3 in dopaminergic neurons. Although the increased amplitude of body-bends phenotype is not rescued by expressing *npr-3* specifically in DVA neurons, however, there seems to be a decrease in the mean amplitude of body-bends in the rescuing line. These data indicate that dopaminergic neurons are involved in regulating npr-3 dependent increase in amplitude of body-bends. (B) The data shows that mutants in *cat-2* display an increased amplitude of body-bends, however the defect is not as severe as that seen in *npr-3 mutants*. Statistical significance in A and B was determined using One-way ANOVA with the Tukey test and represented by “ns” for not significant, “**”, “***’’ or “****” for p-values less than 0.01, 0.001, or 0.0001, respectively

Together, these findings highlight the role of the neuropeptide FLP-15 functioning through the NPR-3 receptor to regulate normal body-bends. We next wanted to test other possible pathways through which FLP-15 may function.

### FLP-15 regulates the expression of NLP-12 to maintain the amplitude of body-bends

Although previous studies have extensively explored the mechanisms behind the sinusoidal wave patterns in *C. elegans* (Chalfie *et al*, 1985; Shingai *et al*, 2013; Wen *et al*, 2012; White *et al*., 1976), the role of neuropeptides in this phenomenon has largely been unaddressed. A recent study by Ramachandran et al. highlighted that overexpressing the neuropeptide NLP-12, specifically expressed in the DVA neuron, results in deeper body-bends in *C. elegans*, suggesting a significant role for neuropeptides in modulating this behaviour (Ramachandran *et al*., 2021).

Given that *flp-15* mutant animals exhibited a similar increase in the amplitude of body-bends, we decided to investigate the expression of NLP-12 in both *flp-15* and *npr-3* mutants using qPCR. The rationale was to see if changes in NLP-12 expression levels could explain the observed phenotype. Our data revealed that NLP-12 was significantly higher in *flp-15* mutants when compared to WT animals. In contrast, *npr-3* mutants showed only a slight increase in NLP-12 gene expression (Figure 5A). Based on these results, we hypothesized that loss of *flp-15* and subsequent increase in NLP-12 expression leads to an increase in the amplitude of body-bends. To test this hypothesis, we conducted behavioural experiments of *nlp-12, and nlp-12; flp-15* mutant *C. elegans*. For the hypothesis to be true we anticipated that mutating the *nlp-12* gene in *flp-15* mutant *C. elegans* would mitigate the increased amplitude defect, restoring the amplitude of body-bends closer to WT levels. Our results showed that *nlp-12* mutants *C. elegans* show a small but significant decrease in the amplitude of body-bends when compared to the wild type animals (Figure 5B). Further, we saw that in *nlp-12; flp-15* double mutants, the amplitude of body-bends was significantly reduced in comparison to *flp-15* mutants and was non-significant from the amplitude of body-bends of WT animals and *nlp-12* mutant *C. elegans* (Figure 5B). These results suggest that the increased amplitude of body-bends observed in *flp-15* mutants are due to the overexpression of NLP-12. The slight overexpression in *npr-3* mutants indicates that while NPR-3 might contribute to this phenotype, we cannot exclude the possibility that other FLP-15 receptors might also be involved in this process.

**Figure 5:**
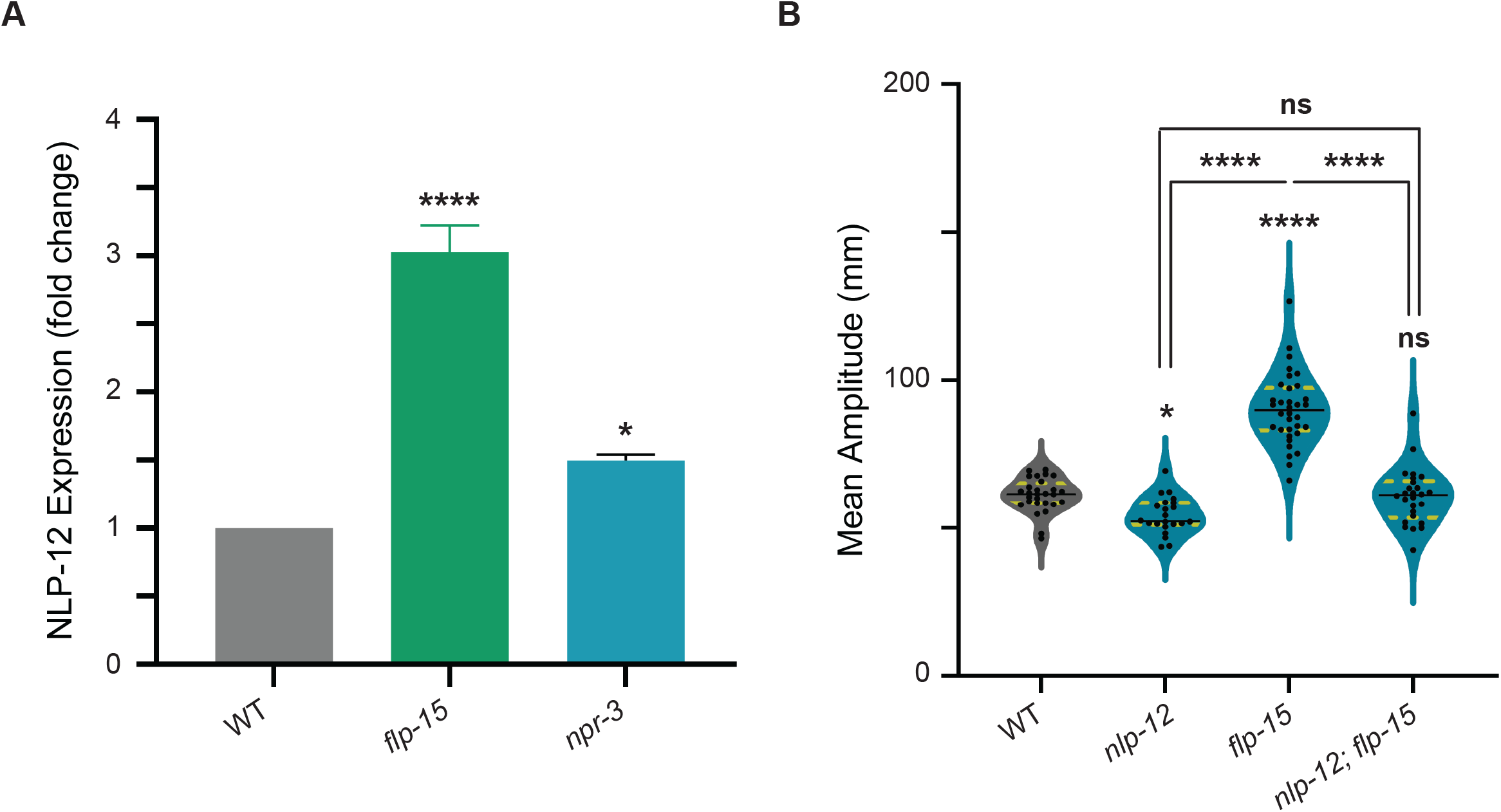
Mutants in *flp-15* show increased *nlp-12* levels. (A) qPCR data shows that *flp-15* mutants show an approximately three-fold increase in the expression of *nlp-12*. However, there is only a 1.5-fold increase of *nlp-12* in *npr-3* mutants. Each bar apart from WT represents the average fold change in *nlp-12* expression in the corresponding mutant background. The error bars represented SEM. (B) Quantification of mean amplitude in *nlp-12* mutants showed that there was a significant decrease in the amplitude of body bend during locomotion. Additionally, introducing *flp-15* mutation in the *nlp-12* mutant background leads to a decrease in the mean amplitude of body bend. Statistical significance in A and B was determined using One-way ANOVA with the Tukey test and represented by “ns” for not significant, “*” or “****” for p-values less than 0.05, or 0.0001 respectively.

Taken together, our work identifies FLP-15 as a novel regulator of the amplitude of body-bends in *C. elegans*. Further, we show that FLP-15 may function through NPR-3 in dopaminergic neurons to regulate the amplitude of body-bends. Finally, our data also suggests that FLP-15 may function via DVA, possibly through other receptor/s to modulate NLP-12 levels that also appears to affect body-bends in *C. elegans* (Illustrated in Figure 6).

**Figure 6:**
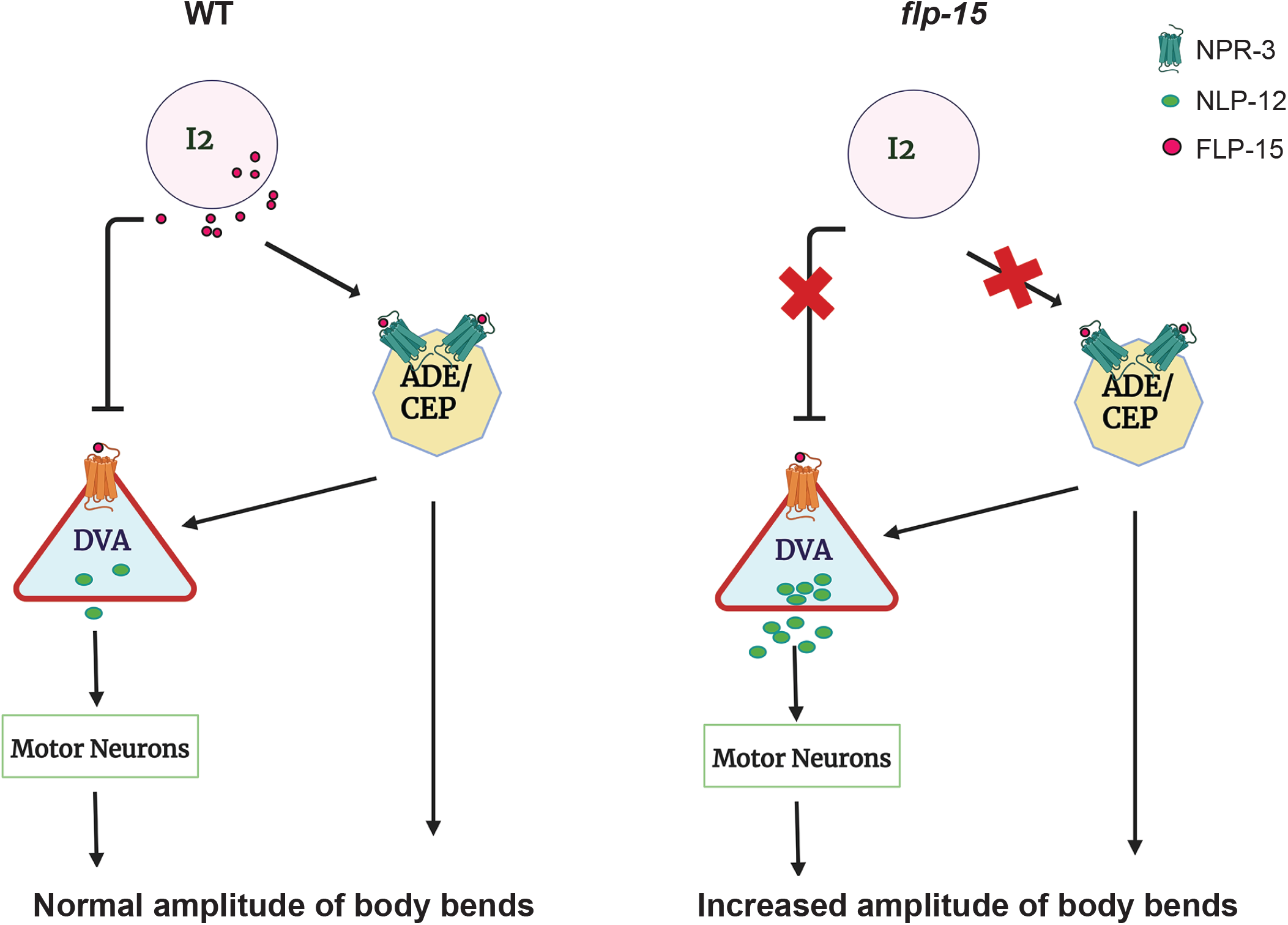
Model indicating how FLP-15 may regulate the amplitude of body-bends. Our data indicates that FLP-15 may function through dopaminergic neurons as well as regulate NLP-12 levels to allow for maintaining the amplitude of body bends in WT *C. elegans*.

## Discussion

*Caenorhabditis elegans* locomotion is characterized by distinct patterns, including forward movement, reversals, and omega turns (Gray *et al*, 2005). Despite numerous studies investigating the neural circuitry controlling these locomotory behaviours, the underlying mechanisms and molecular players remain largely unknown. Most research has focused on the mechanisms governing the frequency of reversals, leaving a gap in understanding aspects of body-bends and the mechanisms underlying them (Bhardwaj *et al*, 2020; Cohen *et al*, 2009; Gray *et al*., 2005; Piggott *et al*, 2011).

Neuropeptides have emerged as crucial regulatory molecules modulating specific locomotor characteristics. They play a significant role in fine-tuning the locomotory circuit in *C. elegans*, influencing not only the frequency of reversals but also other aspects of movement. A study by Hu et al. explored the neuropeptide NLP-12 and its role in a feedback loop that regulates cholinergic transmission at the NMJ. They found that NLP-12 is secreted from stretch-sensitive DVA neurons during muscle contraction, which is stimulated by increased acetylcholine transmission (Hu *et al*., 2011). NLP-12, in turn, activates its receptor CKR-2, further enhancing cholinergic transmission at the NMJ (Bhattacharya *et al*., 2014; Hu *et al*., 2011; Ramachandran *et al*., 2021). This feedback loop underscores the complex interplay between neuropeptides and neurotransmitters in modulating motor functions and maintaining the E/I balance. The FLP-18 neuropeptide has also been implicated in modulating overexcitation in *acr-2(gf)* mutant *C. elegans*.

These mutants carry a gain-of-function mutation in the nicotinic acetylcholine receptor ACR-2 subunit, which leads to overexcitation and a reduction in GABA-induced inhibition, disrupting the E/I balance. The study revealed that the *acr-2(gf)* mutation upregulates FLP-18, which helps restore balance by activating its receptors NPR-1 and NPR-5. These finding highlights the role of FLP-18 in counteracting the effects of overexcitation and maintaining homeostasis in neural circuits (Jospin *et al*., 2009; Stawicki *et al*., 2013). These findings suggest that neuropeptidergic signalling is essential for the precise control of *C. elegans* locomotion Reorientations, including reversals and omega turns, are crucial for food-searching behaviours of *C. elegans*. Bhardwaj et al. found that the length of reversals influences the turning angle during reorientation, with shorter reversals resulting in smaller angles and longer reversals leading to larger angles and omega turns (Bhardwaj *et al*., 2018). In that study, FLP-18 was identified as a key regulator of reversal length. This study builds on the previous work by showing that *flp-15* mutants also have significantly longer reversals compared to WT animals, indicating FLP-15’s role in maintaining normal reversal lengths. We also found that *npr-3* mutants displayed increased reversal lengths, suggesting that FLP-15 and NPR-3 function in the same pathway.

Furthermore, we observed that *flp-15* mutants exhibited abnormal “floral” patterns during reversals, unlike the straight reversals of WT animals. This pattern, which was not as pronounced in *npr-3* mutants, suggests a disruption in normal movement behaviour and efficient foraging. We hypothesized that increased body-bend amplitude in *flp-15* mutants possibly caused this defect, which was confirmed through quantification of the amplitude of body-bends in *flp-15* and *npr-3* mutants. Rescue experiments partially restored normal body-bend amplitude and mitigated the floral pattern. Our qPCR analysis revealed that NLP-12 was significantly overexpressed in *flp-15* mutants, suggesting that this overexpression contributes to the increased body-bend amplitude.

These findings indicate that FLP-15 and NPR-3 are crucial in regulating both the amplitude and directionality of body-bends, highlighting the complex neuropeptidergic modulation of locomotion in *C. elegans*.

Taken together, our data suggests that FLP-15, released from I2 neuron, functions through NPR-3 on dopaminergic neurons to regulate the length of reversals which may possibly function downstream through the release of dopamine. We also show that amplitude of body-bends which further controls the direction of reversal is regulated partially through NPR-3. Further, FLP-15 negatively regulates the expression of NLP-12 to maintain the amplitude of body-bend during locomotion (Illustrated in Figure 6). This highlights a potential mechanism where neuropeptide regulation plays a crucial role in controlling amplitude of body-bends in *C. elegans*, providing new insights into the complex neuropeptidergic modulation of locomotion.

## Acknowledgements

Some strains were provided by CGC, which is funded by NIH Office of Research Infrastructure Programs (P40 OD010440). All illustrations were created in BioRender. We thank Faraz Abbas, Imra Aftab and Velu Saravanan for help with experiments. We also thank Palagiri Suresh for routine help and the members of Kavita Babu’s lab for suggestions and critique on the manuscript.

## Author Contributions

USB: Design and execution of experiments, data analyses, manuscript writing/editing, SH: Execution of experiments, data analyses and manuscript editing, SS: Outcrossing and behavioural screen, NT: Molecular biology experiments, and KB: Supervision, funding acquisition and manuscript writing/editing.

