## Supplementary material for "FLP-15 modulates the amplitude of body bends during locomotion in *Caenorhabditis elegans*"

**Supplementary Information:**

**Table S1 (List of strains)**

| Strain Name | Mutants | CGC Strain | Outcross Status | Figure No. |
|---|---|---|---|---|
| BAB6015 | *flp-15 (*gk1186) | VC2504 | Outcrossed 3X | 1-5 |
| BAB6001 | *npr-3 (ind6001)* | This study | Outcrossed 3X | 3-5 |
| BAB6037 | *flp-15; npr-3* | This study | | 35 |
| BAB6038 | *cat-2* (e1112) | CB1112 | Outcrossed 3X | 4 |
| BAB919 | *nlp-12* (ok335) | RB607 | Outcrossed 3X | 5 |
| BAB6039 | *flp-12; cat-2* | This study | | 6 |
| BAB6040 | *npr-3; cat-2* | This study | | 6 |
| BAB6041 | *nlp-12; flp-15* | This study | | 5 |
| BAB6002 | IndEx6002 *[Pflp-15::*FLP-15cDNA *:: T2A :: GFP ; flp-15]* | This study | | 1,2 |
| BAB6005 | IndEx6005 *[Pnpr-3::*NPR-3gDNA*::GFP; npr-3]* | This study | | 4 |
| BAB6009 | IndEx6009 [P*flp-15*::FLP-15cDNA::T2A::GFP; *npr-3*] (FLP-15 O/E in *npr-3* mutants) | This study | | 3 |
| BAB6010 | IndEx6010 [P*dat-1*::NPR-3gDNA::mCherry; *npr-3*] | This study | | 4 |
| BAB6011 | IndEx6006 [P*nlp-12*::NPR-3gDNA::mCherry; npr-3] | This study | | 4 |
| BAB6012 | IndEx6006 [Pgur-3::FLP-15cDNA :: T2A :: GFP ; *flp-15*] | This study | | 2 |

**Table S2 (List of primers)**

| Genotyping, cloning and qPCR primers | | | |
|---|---|---|---|
| **Gene** | **Sequence (5'-3')** | **Primer Name** | **Comments** |
| *flp-15* | CAGTGAGTCCGAAATTTTAAACCTGA | USB69 | External Forward |
| | AGCATTCTAAAATTGCAGATTACGAC | USB70 | External Reverse |
| | ACTGTGCTGCATGTGTTTCTATG | USB71 | Internal Reverse |
| *npr-3* | GAGAAGGCTCGCTGACATTAATG | USB114 | External Forward |
| | CGATAGTCGTTTTCAGCAATTGTG | USB115 | External Reverse |
| | GAAACGTGCCGATTCGAATGTTG | USB116 | Internal Reverse |
| *nlp-12* | CAATTGCCTCCGACAAAATGC | USB150 | External Forward |
| | CTAACATGTTGACCATACTCTGTG | USB1511 | External Reverse |
| | GATGCGTTTAATTTTGCCATTTG | USB152 | Internal Reverse |
| P*flp-15* | ACACAAGCTTCAATCAATATAGCCGGAAATGCCA | HindIII | UB13 |
| | ACCAGGATCCGTATGTGGGAGACCTTCTTCCA | BamHI | UB14 |
| *flp-15* cDNA | ACCAGGATCCTGGAAGAAGGTCTCCCACATAC | BamHI | UB15 |
| | CACAGGGCCCATTTCATAGAAACACATGCAGCAC | XmaI | UB16 |
| P*gur-3* | ATATGCATGCACATCTCATTGGTTGATTTGATC | SphI | UB25 |
| | CTATGGATCCGAAAACGTGACTGACTAACAC | BamHI | UB26 |

| | | | |
|---|---|---|---|
| P*npr-3* | TGTGAAGCTTCTTAATACTGTCCGTCTGCAAG | HindIII | UB33 |
| | CACACCCGGGTCAAAAATTCCAGAAGAGGAGAAC | XmaI | UB34 |
| *npr-3* gDNA | TCATCCCGGGATCCAAAATGGAGGGTGGTC | XmaI | UB35 |
| | ACACGGTACCTCTAACAACCCGGTAGAATCATC | KpnI | UB36 |
| P*dat-1* | ATCTGCATGCTGCCACCGATTTGTACAAATG | SphI | UB43 |
| | CTATGGATCCGGCTAAAAATTGTTGAGATTCGAG | BamHI | UB44 |
| *nlp-12* | GTATTCGTGGAGGTTTTTGCAAC | RT-PCR Forward primer | |
| | TATTGCCATTCCCAGATTGGCTC | RT-PCR Reverse primer | |

**Table S3 (List of plasmids)**

| Plasmid | Construct | Backbone |
|---|---|---|
| *pBAB6001* | *Pflp-15*::FLP-15cDNA::T2A::GFP | pPD95.75 |
| *pBAB6009* | *Pdat-1*::NPR-3gDNA::mCherry | pPD49.26 |
| *pBAB60010* | *Pnlp-12*::NPR-3gDNA::mCherry | pPD49.26 |
| *pBAB6002S* | *Pgur-3*::FLP-15cDNA :: T2A :: GFP | pPD95.75 |
